# livn: A testbed for learning to interact with in vitro neural networks

**DOI:** 10.64898/2025.12.16.694706

**Authors:** Frithjof Gressmann, Ivan Georgiev Raikov, Hau Ngoc Pham, Evan Coats, Ivan Soltesz, Lawrence Rauchwerger

## Abstract

The investigation of cultured biological neural networks is a critical frontier in neuroscience with profound implications for the advancement of brain-machine interfaces, treatments of neurological diseases, and fundamental insights into neural computation and cognition. Advances in induced pluripotent stem cell (iPSC) technology and machine learning are converging to enable novel approaches to interrogating neural circuits *in vitro*. However, progress in this emerging field is hampered by the technical challenges and resource-intensive nature of acquiring datasets suitable for machine learning. Experimental recordings typically do not allow for interactive learning and generally lack ground-truth information that would enable rigorous algorithm development and validation. To overcome the limitations of experimental setups, simulated biophysical models of neurons and neuronal networks can serve as crucial accelerators. They allow for more controlled and systematic exploration than currently possible with living cultures and serve as interpretable intermediaries between abstract computational theory and complex biological reality. Using this approach, we introduce *livn*: an open source interactive simulation environment for learning to control in vitro neural networks. *livn* generates synthetic neural data with ground truth at scale, enabling the development and testing of ML models in interactive settings that mimic experimental platforms. We describe benchmark tasks that challenge ML models to exploit simulated neural dynamics and release generated synthetic datasets that mimic in vitro systems. By providing an open, extensible platform for developing and benchmarking machine learning models, *livn* aims to accelerate progress in both ML-driven understanding and engineering of in vitro neural systems and fundamental understanding of computation in biological neural networks.

## 1 Introduction

Cultured neural circuits represent a fascinating frontier in computing: living neural networks that self-organize, adapt, and process information through mechanisms refined by evolution. Unlike silicon-based neuromorphic systems or artificial neural networks, these cultures embody the flexible and adaptive nature of biological neural activity, including non-linear dynamics, metaplasticity, neuromodulation, and cell-type diversity that engineering only approximates (Finger, 1994).

These unique properties, combined with recent technological advances, now position cultured neural networks as tractable experimental systems for computational investigations. Advances in stem-cell technology enable the creation of biological neural networks in vitro, offering novel experimental platforms for the exploration of neural systems (Smirnova et al., 2023). Furthermore, increasingly sophisticated machine learning approaches now enable effective data-driven modeling and control strategies for observed neural dynamics (Richards et al., 2019). These developments create opportunities for machine learning models that can learn to interact with bio-engineered living neural networks to decode their activity and contribute to the understanding and engineering of neural processing systems inspired by the brain (Gressmann et al.).

In addition to neuroscientific advances, improved ML-driven control of neural systems has a wide variety of important applications, including the brain-computer interfaces (He et al., 2020), biorobots (Chen et al., 2023), and medical treatments for neural diseases such as Alzheimer’s disease and schizophrenia (Mossink et al., 2021). More broadly, uncovering the principles that make biological neural computation efficient and robust could inform the development of next-generation AI hardware and systems (Gressmann et al.).

However, the computational potential of these living systems remains largely unexplored, due to the lack of systematic frameworks for characterizing their dynamics in terms relevant to computational theory. Experiments with living neural networks remain challenging as they have to contend with a nascent technology stack, crude input-output interfaces, and, fundamentally, an insufficient understanding of the underlying neural mechanisms. Although recording technology and data collection have progressed significantly in recent decades, laboratory experiments remain costly and require hard to come by expertise. For machine learning practitioners in particular, the available neural recording datasets come with crucial limitations that hinder progress in the field. Specifically, most experimental setups provide only crude estimates of critical system parameters–neuron location, connectivity, and synaptic weights–when obtainable at all. This makes it difficult to verify data-driven inferences about the neural dynamics that separate signal from spurious recording artifacts. Moreover, available neural recording datasets are static and preclude interactive input-response feedback loops, thus hindering validation of ML model predictions in closed-loop settings.

Biophysical computational models of neurons and neuronal networks can help in sidestepping experimental limitations by allowing for a more controlled and systematic exploration than currently possible with living cultures. Moreover, biophysical models can serve as interpretable intermediaries between abstract computational theory and complex biological reality. Notably, data-driven systematic characterization of neural dynamics aided by biological models could not only advance neuroscientific understanding, but also lead to novel brain-inspired computing architectures that harness biological principles.

To this end, we develop *livn*^1^ – an interactive, scalable data generator and learning environment that allows machine learning researchers to utilize biophysical neuronal network models to develop and test approaches to **l**earn to interact with **i**n **v**itro neural **n**etworks. *livn* leverages biophysical models to generate high-fidelity neural data at scale, making it possible to obtain ground truth and simulate system responses interactively in closed-loop settings. By *ground truth* we refer to complete access to model variables and parameters during simulation, not to absolute biological ground truth, as all models are approximations of biological reality.

Specifically, in this work, we make the following contributions.

- **We describe an emerging interdisciplinary effort to engineer and control stem cell-derived living neural networks in vitro**. We observe that the lack of ground truth and the static nature of available lab recording datasets make it hard to validate machine learning strategies before their costly deployment in real-world lab experiments. To address this, we propose synthetic data generation from neural simulation.
- **We develop the *livn* environment, a framework for ML-driven neural simulation and control**. Leveraging large-scale neural simulation, *livn* allows for data generation with experimental characterization and control that is comparable with contemporary in vitro lab platforms. We open source *livn* along with generated datasets from six in vitro systems of varying size, complexity and kind. This allows machine learning practitioners to develop and test novel ML approaches against simulated experimental environments.
- **We describe possible machine learning tasks for open- and closed-loop experiments**. While *livn* does not implement specific, strictly standardized benchmarks, we provide a general problem formulation that challenges a given machine learning model to solve a task by *exploiting* the in vitro neural system. This leaves room for practitioners to pick relevant tasks at a suitable level of difficulty, allowing for gradual increasing complexity to drive methodological progress.

*livn* is released under the permissive MIT license and designed to be extensible at multiple levels to reflect the great variety of possible platforms and systems used in experimental laboratories. Users are free to implement custom neural dynamics, adjust default parameters, connectivity, and architectures, and introduce or extend input-output mechanisms and devices.

Although *livn* includes a set of tuned standard models, we encourage users to refine and contribute their own models to closely reflect a given experimental platform. Over time, we hope that *livn* can serve the community by integrating expertise in neural simulation, state-of-the-art experimental techniques, and machine learning to develop and rigorously benchmark models before their ultimate validation in lab experiments.

While direct validation against experimental electrophysiological recordings is beyond the scope of this initial release, future work will systematically compare simulated response statistics (firing rates, burst properties, correlation structures) to published in vitro culture data. We anticipate such validation will guide refinement of model parameters and may reveal which biological mechanisms are most critical for capturing experimental phenomenology.

## 2 Motivation

The breakthrough development of induced pluripotent stem cells (iPSC) has transformed the experimental capabilities for biomedical and bioengineering research (Ye et al., 2013; Takahashi et al., 2007; Kegeles et al., 2020; Mossink et al., 2021). In particular, “organoid” technology now allows for the routine generation of increasingly sophisticated 3D cell cultures outside of their usual biological context *in vitro* (see Zhao et al. (2022) for a comprehensive review).

Using 3D organotypic culture techniques, researchers can generate functional neuronal networks that, while much simpler than intact brain structures, exhibit complex activity patterns and network dynamics. The in vitro neuronal networks (IVN) can be interfaced bidirectionally through multi-electrode arrays (MEAs) that record extracellular field potentials and deliver targeted electrical stimulation to specific neuronal populations (Chen et al., 2017). Furthermore, neurons can be transfected with viral vectors expressing photoreceptor proteins (e.g. channelrhodopsin, halorhodopsin) to enable millisecond-precision control of neural circuitry through light stimuli (Fenno et al., 2011). By integrating these electrophysiological and optogenetic techniques, investigators simultaneously manipulate and monitor spatio-temporal activity patterns throughout the IVN, providing access to the emergent dynamics of the neural circuits.

Building on these advances, it is possible to perform closed-loop experiments in which the measured neural activity affects an environmental state that is subsequently fed back into the IVN (Kagan et al., 2023). For example, Kagan et al. (2022) demonstrated that the activity feedback of in vitro neurons can be used to perform basic video game play. In this setup, a predefined encoding and decoding mechanism translates between the IVN activity and a simulated ‘pong’ game world, allowing the in vitro systems to adapt its response based on the environmental feedback. Notably, general machine learning techniques are increasingly successful in inferring sophisticated decoding and control strategies from neural data. For instance, Li et al. (2024) used a reinforcement learning (RL) algorithm to learn to motor-control a C. elegans nematode. The policy was trained offline on pre-recorded data to optimize the light stimulus used for optogenetic stimulation.

However, in practice, continued development and testing of machine learning approaches for lab experiments remain challenging and bottlenecked for several reasons. First, experimental data collection from lab environment is difficult to automate and scale, making it difficult to generate sufficient training data for modern deep learning architectures. For instance, Li et al. (2024) noted that it was infeasible to collect thousands of hours of recordings that are typically used by deep RL methods trained with simulators. Secondly, when working directly with a living system, it is challenging to disentangle whether optimization issues stem from the machine learning model or environmental factors. In the case of C. elegans training, without the ability to diagnose sudden performance drops, the authors had to rely on ensembles of agents to reduce variance and achieve training stability (Li et al., 2024). Finally, and perhaps most crucially, data from lab experiments can by definition only offer the observable system state making it harder to validate hypothesis about the system. For example, while the Pong gameplay experiment (Kagan et al., 2022) demonstrated the system’s ability to adapt in response to stimuli, it revealed little about the underlying neural mechanisms involved in the process. Due to recording limitations, it is, for example, not possible to track synaptic weight changes over the course of the stimulation. More generally, the ever evolving nature and lack of ground truth of living neural systems means that valuable validation techniques such as ablation studies or exact experiment replications are not available, hampering ML innovation in the field.

Fortunately, given the ever-increasing computing resources, synthetic data generation through neural simulation is now becoming increasingly feasible (de Melo et al., 2022). Notably, the NEURON simulation engine (Hines et al., 2022) that is used by projects such as NetPyNE (Dura-Bernal et al., 2019) and BioNet (Gratiy et al., 2018; Dai et al., 2020) enables simulation of detailed, multi-compartment neuronal models at super-computing scale. However, existing solutions are not geared for data generation for machine learning and specifically lack comprehensive models of in vitro cultures and platforms. Motivated by these limitations, we design the *livn* framework that enables machine learning practitioners to develop and validate ML techniques for in emerging in vitro systems.

## 3 The *livn* environment

*livn* builds on the long-standing efforts towards high-fidelity simulations of biological neural systems to allow for synthetic data generation for machine learning at scale. This includes simulation of neural dynamics as well as input-output transformations that model experimental setups such as multi-electrode arrays and optical stimulation. To support the simulation of millions of neurons on high-performance computing infrastructure, *livn* leverages the NEURON simulator (Hines et al., 2022) and the neuroh5 library (Raikov et al., 2021) to provide MPI-based scaling for detailed large-scale biophysical models with topographical connectivity (Upadhyay et al., 2023). Figure 1 and Table 1 provide an overview of the framework.

**Table 1.** Overview of the default *livn* systems and their the generated datasets. Each data sample consists of a simulation of 5 seconds of physical time under varying feature inputs.

| Cell Population | Name | Neurons (exc./inh.) | Dataset samples (train/test) |
| --- | --- | --- | --- |
| EXC-INH | S1 | 10 (7/3) | 50k / 1k |
| | S2 | $10^2$ (80/20) | 50k / 1k |
| | S3 | $10^3$ (800/200) | 5k / 500 |
| | S4 | $10^4$ (8000/2000) | 500 / 50 |
| Hippocampal | C5 | $\approx 10^5$ (15 cell types) | single |
| | C6 | $\approx 3.5 \times 10^6$ (15 cell types) | single |

**Figure 1.**
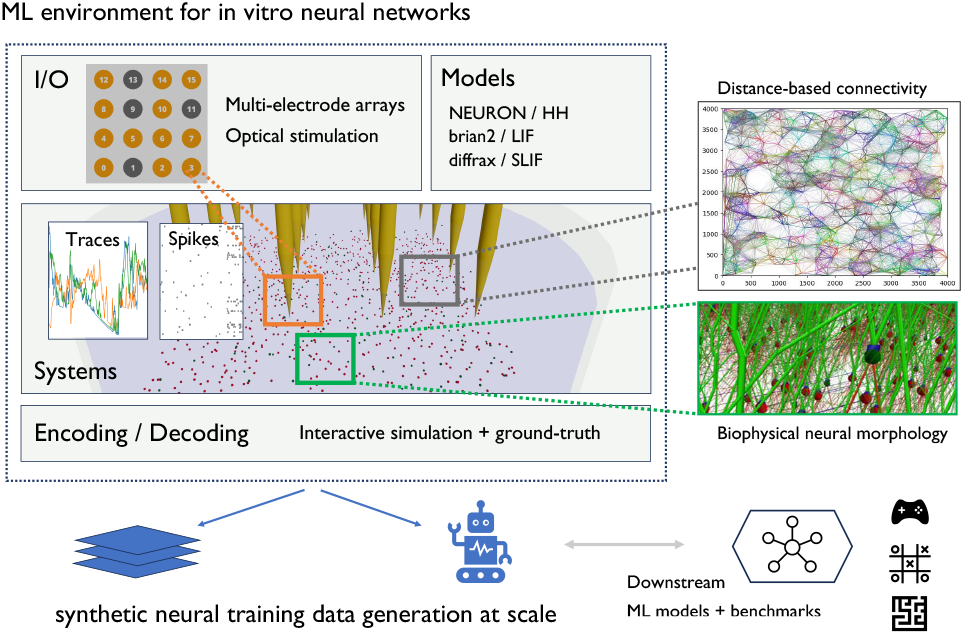
Overview of the *livn* framework that aims to enable synthetic neural data generation for machine learning with in vitro neural networks. Large-scale biophysical models are supported via NEURON (Hines et al., 2022) while simpler models such as leaky-integrateand fire (LIF) and Izhikevich (Izhikevich, 2003) can be implemented using brian2 (Stimberg et al., 2019). Furthermore, *livn* supports differentiable dynamics models such as Stochastic Spiking Neural Networks (Holberg & Salvi, 2024) implemented in diffrax (Kidger, 2022). The environment supports dataset generation as well as interactive learning, providing integration for downstream machine learning benchmarks via the Gymnasium standard interface for RL environments (Towers et al., 2024).

### 3.1 Simulator

At a basic level, *livn* users can configure preexisting or custom environments that specify the system (neuron locations, connectivity, etc.), the model (dynamics, synapses, etc.), as well as the IO (electrodes, optical stimulation, etc.). After initialization, the environment allows to simulate the system for a given stimulus and specified amount of physical time. Both spiking and cell voltage traces can be recorded as a simulated activity response. The user is free to continue the simulation with an altered stimulus at any time, allowing for the implementation of continuous simulation and closed-loop setups.

#### System generation

Unlike their artificial counterparts, biological neurons feature rich morphologies with thousands of synaptic connections for a single cell. To generate systems, users can specify the number of neurons, system size, and cell morphology using the SWC standard (Halavi et al., 2008) to generate realistic three-dimensional neural architectures that mimic the spatial organization observed in biological neural circuits. *livn* includes six predefined systems with varying neuron counts described in Table 1. Specifically, livn includes four EXC-INH systems that consist of 80%/20% excitatory and inhibitory cells, respectively. This mimics stem-cell-derived mixed cultures where directed differentiation creates excitatory and inhibitory neuronal subtypes of known ratio. Secondly, livn includes two hippocampal systems that model the CA1 region of a rodent hippocampus. This can be used to model in vitro brain slices from cortical organoids or mice.

#### Models

Given the cell location and connectivity of the system, users can choose from a variety of predefined models or implement custom neural dynamics. In particular, *livn* supports three simulation engine backends to express models of neural dynamics, namely NEURON (Hines et al., 2022), diffrax (Kidger, 2022) and brian2 (Stimberg et al., 2019). This provides flexibility in the level of model abstraction and complexity, allowing to study the impact of different dynamic models and fidelity for a given system. Notably, diffrax-based models such as Stochastic Spiking Neural Networks (Holberg & Salvi, 2024) allow for exact gradient computation, making it possible to differentiate through simulations end-to-end.

The predefined NEURON models used for synthetic data generation are informed by experimental summary data to produce realistic models of in vitro cultures despite the many experimental gaps. Specifically, the EXC-INH synaptic parameters are tuned such that the neural populations produce the reported mean-firing rates around 3 Hz and 12 Hz for spontaneously active and stimulated populations respectively (Zhang et al., 2023). Furthermore, the two-compartment Pinsky-Rinzel motoneuron models have been constrained using the reported data from electrophysiological experiments by Miles et al., including f-I curves, input resistance, and membrane capacitance. For the hippocampal system, we rely on the quantitative estimate of the cellular and synaptic components by Bezaire & Soltesz (2013). More details can be found in the supplementary material.

#### IO

While *livn* provides full access to the simulated neural dynamics, machine learning models should be designed and trained in a way that is directly applicable to real-world in vitro systems. In particular, training data should mimic the level of experimental characterization and control that is common for contemporary in vitro lab platforms. To account for this, *livn* allows to specify IO-transformations that translate between the neuronal dynamics and the experimental channels that would be stimulated (In : ℝ^channels^ → ℝ^neurons^) or read-out (Out : ℝ^neurons^ → ℝ^channels^) during the experiment. For example, an Out-transformation could drop detected neuron spikes probabilistically based on distance to mimic noisy electrode spike detection of nearby neurons.

*livn* implements transformations to simulate multi-electrode arrays using a point-source model, which computes the amplitude of the signal detected by an electrode based on the distance between the neuron and the electrode. Specifically, we model the electrode induced voltage *v* for each neuron as In^MEA^ : *v* ~ *ρI/*4*πr* where *ρ* is the tissue resistivity, *I* is the electrode current, and *r* is the distance between the neuron and the electrode. Notably, this transformation allows for backpropagation when used with the differentiable diffrax simulations. Similar transformations can be defined for models of optogenetic stimulation, where the induction of the optic current is determined by a light source. Specifically, *livn* integrates with the cleo library (Johnsen et al., 2023) that offers a variety of laser and opsin models that can be used to define suitable IO transformations.

#### Encoding, decoding and pattern generation

Neural activity relies on spatio-temporal coding strategies to represent and process information. To encode data into the system’s input channels and neurons, *livn* includes a library of common rate, spike and temporal coding strategies. This allows to characterize neural responses under diverse spatio-temporal input patterns. For temporal signal processing, we choose a spectral filtering approach that models frequency-selective responses similar to those observed in the auditory system. For spatio-temporal features, we include additional spatial dimensions. We describe the details of the dynamical response characterization (DRC) in the supplementary material.

#### Visualization

To aid development and debugging and allow visualization, *livn* can generate browser-based interactive renderings of systems and their neural morphology using the sharkviewer library (Weaver et al., 2014) and includes a widget to visualize systems within Jupyter notebooks (see Figure 1).

### 3.2 Machine learning problem formulation

*livn* is intended as a general purpose neural simulation framework that can aid the development and validation of machine learning approaches for a range of IVN-related challenges, including learned experimental control, ML-driven post-processing and data analysis, or techniques for simulated-based inference (Cranmer et al., 2020). In practice, the broad objective of learning to interact and control in vitro neural networks will need to be broken down into subproblems and ML benchmarks that suit the experimental capabilities of available systems. Notably, the specific ML benchmark problems are intended as a tool to assess the level of control achieved; given a standard *livn* system and standard ML benchmark, the challenge is learning to control the livn system such that it solves the benchmark. As such, we find that a strict definition of a novel set of machine learning benchmark tasks could be too limiting to reflect the broad range of practically relevant ML problems and available systems. Instead, inspired by the open division of MLPerf (Reddi et al., 2020), we formulate a semantic-level problem description that leaves room to pick the appropriate established ML tasks and implementations (see the illustration in Figure 2).

**Figure 2.**
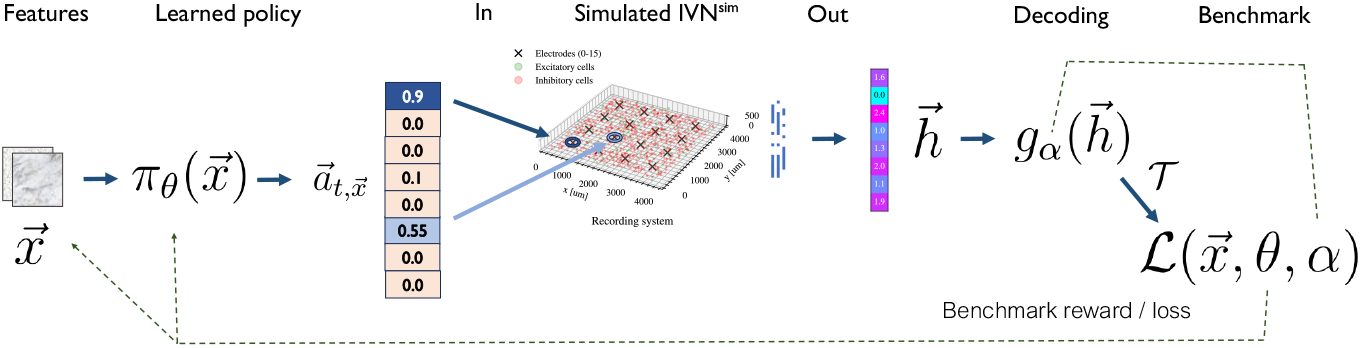
Illustration of the sequential decision problem for interaction with (simulated) in vitro neural networks. The goal is to learn a policy for the interaction with the in vitro system. To assess learning progress, the output of the IVN is fed into a given machine learning benchmark that provides the overall optimization objective (e.g. cumulative reward in a simulated RL environment). As such, the setup challenges the ML model to learn to exploit the IVN to solve the downstream task.

Specifically, we can view the in vitro neural network as a probabilistic function that maps encoded inputs to decoded outputs, much like a conventional stochastic artificial neural network. However, unlike in the artificial case that may allow easy modification of internal parameters, the functional mapping of in vitro systems can only be controlled through a sequence of inputs whose rules are at least partially unknown. In other words, the interaction with an IVN represents a sequential decision problem where decisions over time affect future states and outcomes. Crucially, the goal or objective (e.g. cumulative rewards) of the problem can be provided through existing ML benchmark problems that consume the decoded output of the IVN. For example, we can choose a standard reinforcement learning problem such as the Pong game where the action is the decoded output of the IVN (e.g. the sub-policy in the hierarchical reinforcement learning formulation). More generally, we can specify a *livn* benchmark as a composed sequential decision problem 𝒯○ Out ○ IVN^sim^○In where 𝒯 is an existing ML benchmark and IVN^sim^ the simulated or observed neural system. In this formulation, learning to solve the benchmark implies learning to exploit the dynamics of the IVN in a way that minimizes 𝒯’s objective. Notably, the benchmark solving abilities provide an implicit measure of successful interaction with the IVN. To foster standardized experimentation in this framework, *livn* integrates with the Gymnasium standard interface for RL environments (Towers et al., 2024), making it easy to create suitable problem compositions.

### 3.3 Dataset generator

The above problem formulation motivates and informs the dataset generation (see Table 1). For each system, the generated dataset should contain a set of simulations (i.e. samples drawn from IVN^sim^) that represent the wide range of possible stimulus-response patterns of the system. This means that the spatio-temporal stimuli used during simulation need to representatively cover the input state space to capture relevant dynamic transitions that allow for effective learning. To this end, *livn* supports parallel sample generation from user-defined feature spaces. We generate and publish dataset of samples for each of *livn*’s predefined systems. The datasets can serve as a starting point for the development of machine learning strategies that learn to make sense of neural activity. For instance, *livn*’s documentation includes an example on how to use the pregenerated data as a replay buffer to bootstrap an off-policy RL agent.

## 4 Experiments

This section presents experiments to establish that (1) *livn* enables scalable data generation of complex biological models with (2) realistic and rich dynamics for a wide range of input features. Figure 3 provides a visualization of a generated dataset sample, i.e., the simulated neural dynamics of an in vitro system in response to stimulation.

**Figure 3.**
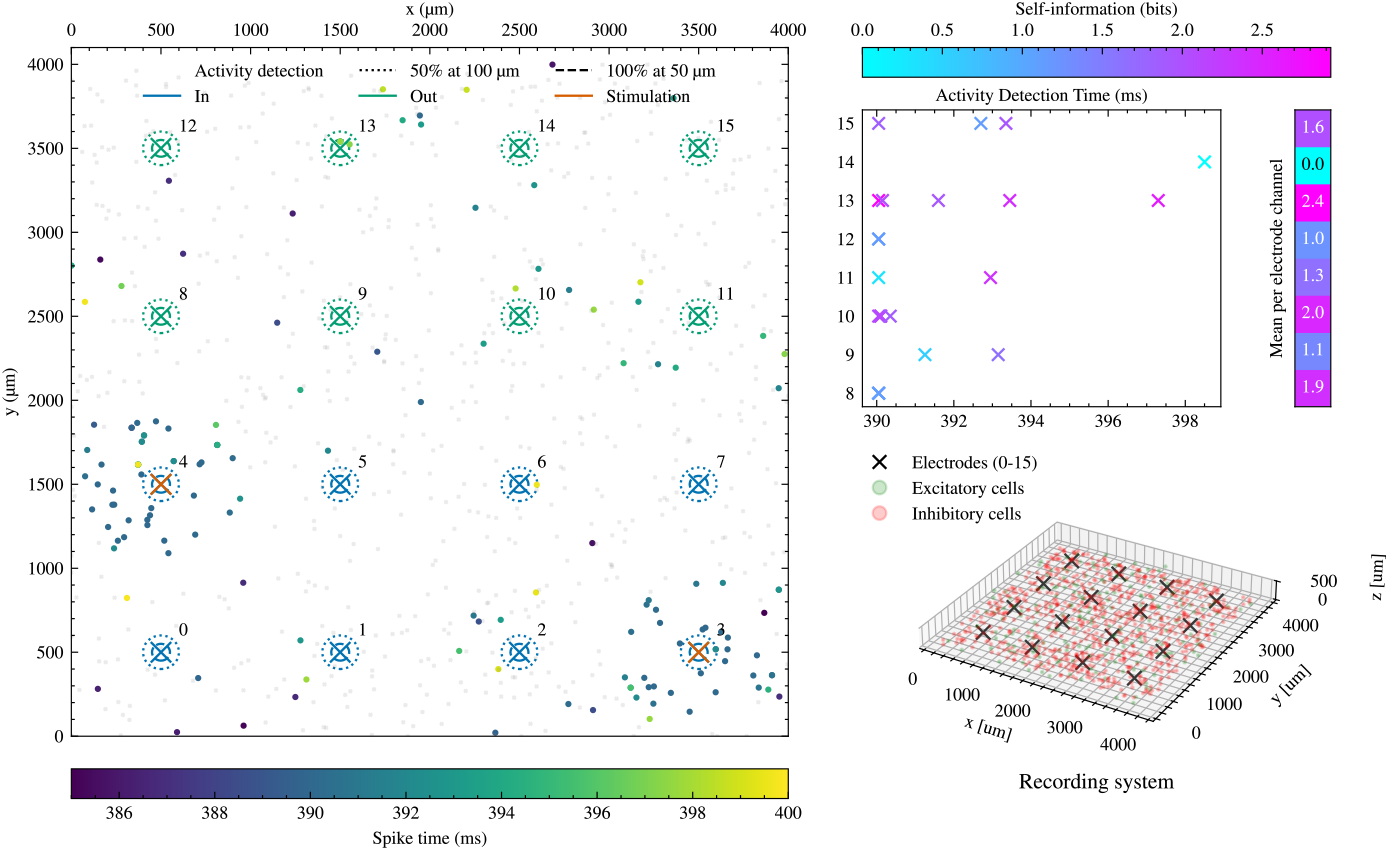
Overview a simulated livn system, illustrating the neural activity and a possible decoding method in response to stimulus. Spontaneous neural activity is recorded for 390 ms, at which point the stimulus is provided at electrodes 3 and 4. The left panel gives a top-down view of the spatial distribution of neurons and their firing activity over time. The top-right panel shows detected activity by the output electrodes 8-15 in the 10 ms following the stimulus. Colors indicate the information content measured relative to the firing activity distribution that was observed before the stimulus.

### 4.1 Scalability

To be able to generate sufficiently large datasets suitable for deep learning, the data sampling needs to be possible at scale. We obtain all simulated results and generated datasets using the NSF leadership-class super-computing systems Frontera (Stanzione et al., 2020; TACC, a) and Vista (TACC, b). Figure 4 presents simulation time measurements with increasing compute resources for both the EXC-INH and Hippocampal systems on Vista and Frontera, respectively. With sufficient compute resources, the EXC-INH simulation step time can be reduced to match the simulated physical time. Furthermore, the smaller EXC-INH S1 and S2 are suitable for use on few-core machines like personal computers and do not require super-computing infrastructure.

**Figure 4.**
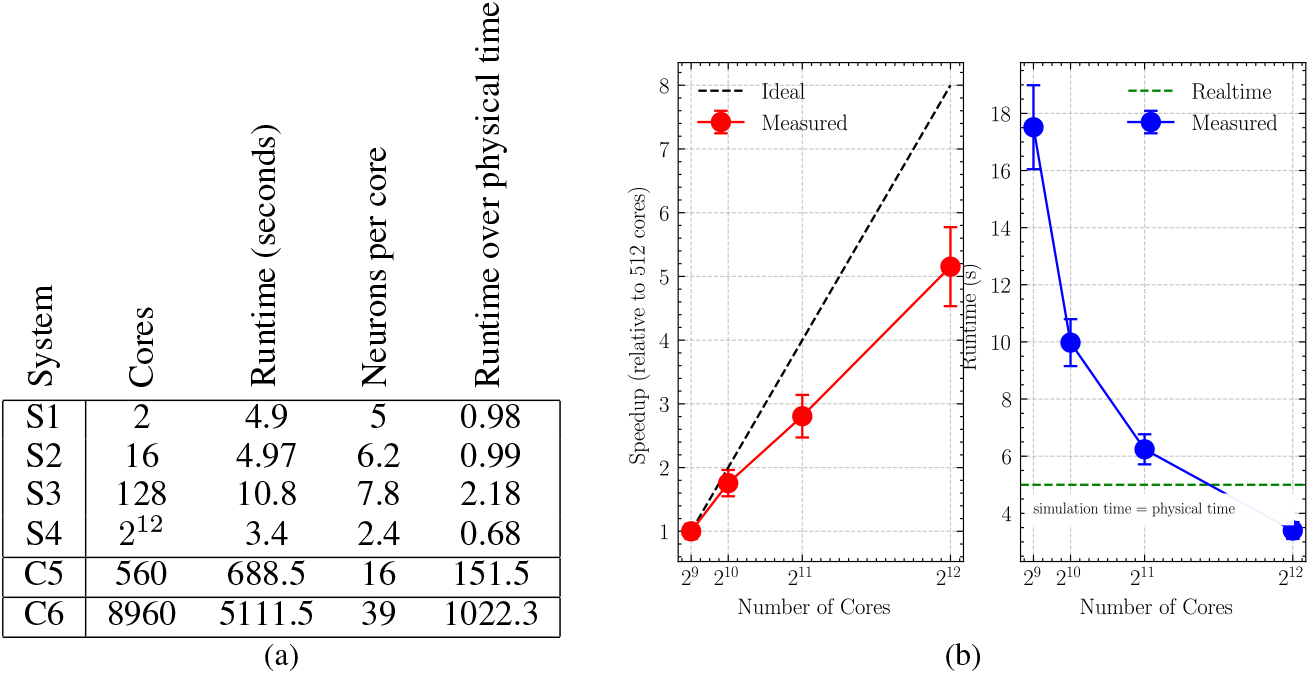
Simulation performance for 5 seconds of simulated physical time of the different *livn* systems using the default NEURON simulator (Hines et al., 2022) models. (a) Scaling of cores with system size and (b) resource scaling of the largest EXC-INH 10k neuron system. Measurements and standard error across 3 trials are obtained on grace-grace nodes of the Vista computing system using NEURON’s pc.step time profiler that measures CPU-time spent integrating equations, checking thresholds, and delivering events. The large-scale hippocampal system C5 and C6 are evaluated on the Frontera computing system.

### 4.2 Stimulus generation and resulting dynamics

To elicit realistic neuronal responses, we generate complex temporal and spatial stimuli that model how neural populations encode sensory inputs across different modalities (see Section 3.1 and Supplementary Material).

Figure 5 visualizes the simulated stimulus-response characteristics of *livn*’s largest C6 system that models hippocampal CA1 dynamics. The metrics confirm a non-pathological response with most cells being active and some showing preferences for specific frequencies in the input signal. The changing network excitability in response to different input frequencies indicates that it should be possible to influence and control the dynamics through input signal modulation. As such, this presents an exciting challenge for machine learning approaches that can uncover effective control policies for complex neural dynamical data.

**Figure 5.**
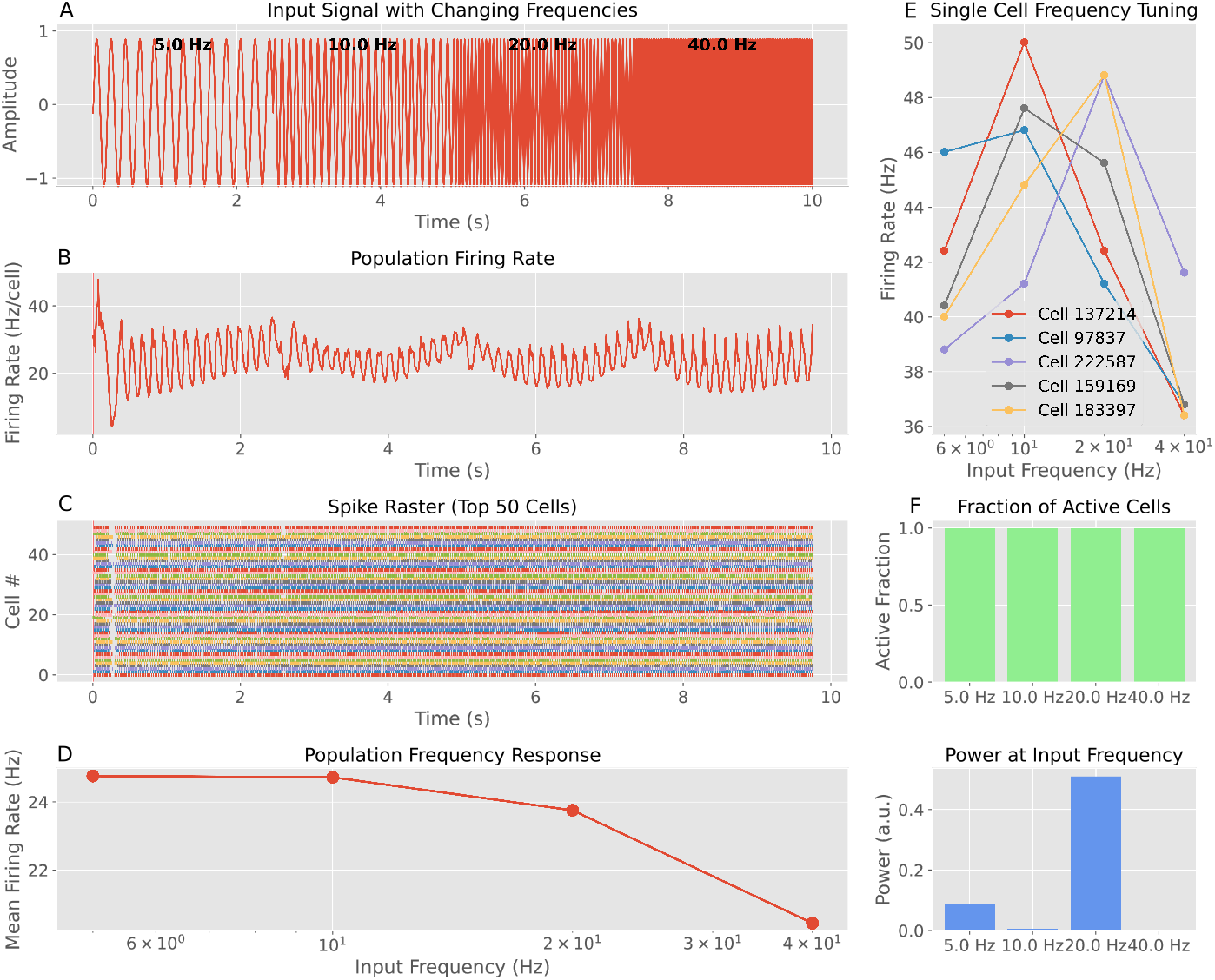
Visualization of the input-output transformation performed by the C6 system (hippocampus CA1 network) on temporal frequency input signal. A) The input signal consists of four segments with different oscillation frequencies (5Hz, 10Hz, 20Hz, 40Hz) simulating distinct temporal patterns. B) The population firing rate shows the collective activity of CA1 neurons, revealing how network excitability changes in response to different input frequencies. C) The spike raster displays activity patterns of the most active neurons, showing cell-specific responses to each frequency segment. D) The frequency tuning curve quantifies the population’s preference for different input frequencies. E) Single cell tuning curves illustrate the heterogeneity in frequency selectivity across individual neurons, with some cells showing clear preference for specific frequencies. F) Additional metrics show the power at input frequency bands and the fraction of active cells per frequency, revealing how network recruitment varies with input characteristics.

## 5 Discussion and Future Work

The introduced generated datasets and data generation capabilities make it possible to obtain neural data with ground truth that would be unobtainable from real-world experiments. However, it is important to note that simulated neural data comes with crucial limitations. Biological reality is too complex to be captured even by the most sophisticated computational models. This means that any models trained on synthetic data will not easily transfer to real-world settings. Given our still limited understanding of neural information processing, it is possible that there are critical properties of neural systems not captured by contemporary biophysical models. Notably, given the relatively recent development of in vitro organoid systems, their experimental characterization is still lacking, as systematic characterization of living neural systems remains notoriously difficult and expensive. For example, the *livn* models currently do not account for synaptic plasticity and rewiring, representing a distinct time in the development of the culture. Thus, it is important that machine learning approaches maintain awareness of the modeling *assumptions* constituted in the synthetic training data. Confirmation of hypotheses must ultimately be attained from real-world data.

For this reason, *livn* only presents a starting point for further development of simulation-driven learning to interact with in vitro systems. While *livn*’s default systems and datasets are informed by the features and capabilities of contemporary in vitro platforms (Zhang et al., 2023; Elliott et al., 2023; Jordan et al., 2024), further improvement and validation against lab experiments remains a crucial priority. Notably, the initial dataset release is only modest in size, aiming to serve as a testbed for exploration and development. We plan to update and release more extensive datasets as ML model requirements and benchmarks are being refined (cp. Section 3.2). One key question is what level of modeling abstraction and simulation fidelity is needed to capture and represent important properties of in vitro systems. With the ability to vary IO, models and information encoding strategies, *livn* can support systematic studies to could help answer these questions. However, a current limitation of the *livn* library stems from the generic nature of the simulated systems that do not yet represent any particular real-world platform.

Our current models do not incorporate several biological mechanisms known to be important in living neural networks:

- Synaptic dynamics: The models represent networks at a fixed developmental timepoint without synaptic plasticity, rewiring, or long-term potentiation/depression. Living cultures exhibit substantial connectivity changes over hours to days that would affect computational properties.
- Neuromodulation: In vitro cultures contain and release neuromodulators (dopamine, serotonin, acetylcholine) that dynamically adjust network excitability and information processing. Our models use fixed synaptic and channel parameters.
- Cellular heterogeneity: While we implement distinct excitatory and inhibitory populations, real cultures exhibit greater cell-type diversity and more complex developmental trajectories than our models capture.
- Biophysical simplifications: Reduced compartmental models (2-compartment for EXC-INH systems) necessarily omit detailed dendritic integration, back-propagating action potentials, and fine-scale morphological effects that may influence network computation.

These limitations mean that machine learning models trained on synthetic data will require adaptation for real experimental platforms. However, the systematic characterization enabled by simulation can guide experimental design and establish computational baselines for comparison. Validation against laboratory experiments remains a crucial priority for future work.

## 6 Conclusion

In this work, we introduce *livn*, a framework that advances ML-driven engineering and control of in vitro neural networks. Through large-scale neural simulation, *livn* can generate synthetic datasets with fine-grained control and ground-truth annotations, thus enabling the development and validation of machine learning models that learn to make sense of the system dynamics. We hope that continued ML progress in this field, guided by feedback from real-world experiments, will help pave the way for increasingly sophisticated computing applications in vitro.

## Ethics statement

Although *livn* is a simulation environment, it is worth noting the potential ethical issues of the research that it seeks to support. Experiments involving living organisms and stem cell technology raise concerns about informed consent, as tissue donors often remain unaware that their cells could be used for iPS cell derivation or neural systems based on their genetic material (Hyun et al., 2020; Kagan et al., 2023). Furthermore, advanced in vitro neural systems might theoretically develop consciousness capable of suffering (Hyun et al., 2020; Goddard et al., 2023; Reardon, 2020), although this remains unlikely near-term (Milford et al., 2023). These considerations need not stop progress, but highlight the importance of developing robust consent protocols and proactive ethical frameworks as the field evolves.

## Acknowledgments

The work was funded by NSF Expedition “Mind in Vitro” award #IIS–2123781. The simulations performed as part of this work used the Frontera supercomputer at the Texas Advanced Computing Center (TACC) through allocation IBN22011, provided through the NSF Petascale Computing Resource Allocation program.

## A Appendix

### A.1 Details of the simulator

*livn* separates the system specification, i.e. the neuron types, morphology, locations, and connectivity, from the possible neural dynamics model (e.g. Hodgkin-Huxley, Leaky-integrate and fire, Izhikevich, etc.). The system is saved in an HDF5-based data format that specifies cell attributes and the connections between population of cells. We take advantage of the neuroh5 library (Raikov et al., 2021) that implements scalable MPI-based construction of large-scale neural network models using the parallel graph partitioning algorithm of Karypis et al. (2003). The user can specify callback functions that construct the specific dynamics and connectivity of the cells while leaving the parallel data movement to the library. A more detailed description of data structures and parallel data movement can be found in Section 2 of Raikov et al. (2021).

#### A.1.1 System generation

To generate the default *livn* systems, we define a rectangular prism that represents the dish with the growing neurons of the experimental platform (see Table 2, cp. example system by Zhang et al. (2023)). We distribute the neuronal somata (cell positions) quasi-uniformly using a dispersion-based optimization within the volume that minimizes intersomatic distances. Volume dimensions and cell densities correspond to those of the rodent hippocampus. This leads to biologically plausible and energy-minimizing cellular arrangements. With the somas in place, we use the stereotyped morphological information of the SWC file (Halavi et al., 2008) to determine the likely locations of the synapses. Specifically, given a synapse density, each segment of the cell’s morphology receives synapses such that the spacing between their locations follows a Poisson distribution. Subsequently, the connectivity is determined as a function of spatial distance between synapses, using axonal extents that are consistent with experimental data. Notably, morphological reconstructions are only used during the generation of network connectivity, not for the dynamical models. Although microscopic differences in connectivity could effect changes in the computational properties of the network, empirical evidence from our simulations seems to suggest that neuronal and synaptic dynamics exert a much greater influence on those properties. The exact configuration and morphology files to reproduce our published system HDF5 files can be found in the code repository at https://github.com/livn-org/livn/tree/main/systems.

**Table 2.**
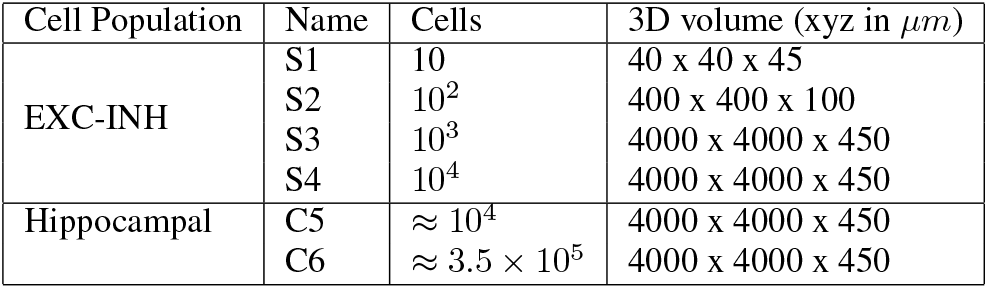
*livn* system volume specification.

| Cell Population | Name | Cells | 3D volume (xyz in $\mu m$) |
| --- | --- | --- | --- |
| EXC-INH | S1 | 10 | 40 x 40 x 45 |
| | S2 | $10^2$ | 400 x 400 x 100 |
| | S3 | $10^3$ | 4000 x 4000 x 450 |
| | S4 | $10^4$ | 4000 x 4000 x 450 |
| Hippocampal | C5 | $\approx 10^4$ | 4000 x 4000 x 450 |
| | C6 | $\approx 3.5 \times 10^5$ | 4000 x 4000 x 450 |

#### A.1.2 Dynamic models

The computational neuroscience literature has proposed a wealth of models for neuronal dynamics, ranging from simpler (stochastic) leaky integrate-and-fire (Gerstner & Kistler, 2002) (LIF) models to more evolved Hodgkin-Huxley type models driven by ionic conductances. *livn* encourages exploration of different models by specifying the cell population and connectivity, but leaving the choice of the exact dynamics model to the user. Out-of-the-box, *livn* includes brian2 implementations of the popular LIF (Gerstner & Kistler, 2002) and Izhikevich (Izhikevich, 2003) dynamics, as well as SLIF (Holberg & Salvi, 2024) that is implemented using diffrax. These point-neuron models trade biophysical realism for speed, allowing simulation of thousands of neurons without supercomputing resources, making it particularly well suited for interactive learning. Conversely, for the dataset generation, we implement more biorealistic, computationally expensive ion-channel models using NEURON which we describe next.

##### NEURON EXC-INH motoneurons

The S1-S4 EXC-INH circuit models represent simplified mixed excitatory-inhibitory cultures with 80/20 ratio consistent with cortical-like differentiation protocols (cp. (Zhang et al., 2023)). While we use Booth et al. (1997) reduced models for computational tractability, these should be understood as generic excitable neuron models rather than biophysically detailed motor neurons.

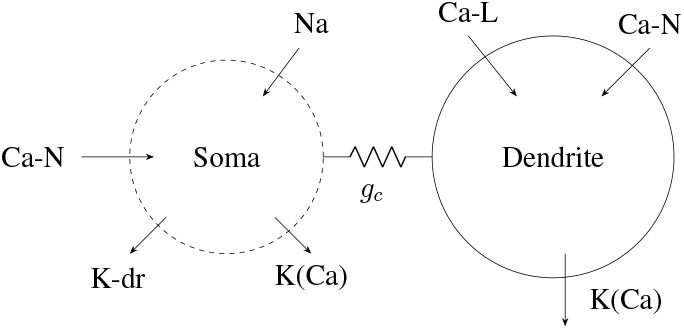

The excitory neuronal model is constrained using data obtained from electrophysiological experiments by (Miles et al.), including f-I curves, input resistance, and membrane capacitance. Furthermore, the synaptic currents are based on experimentally-obtained amplitude and time constant information, while the connectivity is based on known anatomical connectivity distributions. In addition, the inhibitory cells that form 20% of the culture are modeled as parvalbumin-expressing basket cells (PVBC) using a Pinsky-Rinzel model (Pinsky & Rinzel, 1994). Note that while these models do not include full morphological information, the morphology-driven connectivity generation as described above is still being used. This reduces computational complexity while preserving plausible connectivity patterns. Furthermore, the illustrated ion channel complement represents a simplified implementation constrained by available computational resources and experimental data. While based on the (Booth et al., 1997) formalism, the specific channel complement may differ from that of iPSC-derived motor neurons and is better characterized as a generic excitable cell model suitable for capturing basic excitability and adaptation properties.

##### NEURON Hippocampal circuit

The C5 and C6 circuit models represent organoid systems patterned after the CA1 region of the rodent hippocampus. These models extend the foundational full-scale network architecture of Bezaire et al. (2016) by incorporating additional interneuron classes and synaptic dynamics that aim to accurately represent the complex microcircuitry of the hippocampus. The model system consists of 15 cell types as specified in Table 3 and Figure 6. Three additional interneuron-specific classes have been added (IS1, IS2, IS3) that represent functionally specialized inhibitory populations with unique connectivity patterns and laminar distributions. The synaptic transmission mechanisms have been extended by adding NMDA receptor kinetics based on the model developed by Tomko et al. 2021. Beyond the original connectivity matrix developed by Bezaire et al. (2016), the CA1 circuit model incorporates additional connectivity projections derived from the anatomical structure of axonal and dendritic overlap patterns across the hippocampal layers. This extended model architecture allows for investigation of how interneuron diversity and synaptic dynamics contribute to the function and information processing of the CA1 network.

**Table 3.** Cell types and cell count of cortical hippocampal circuit model (C5/C6)

| Cell type | Cell count |
| --- | --- |
| Pyramidal Cell (PYR) | 311500 |
| Parvalbumin-expressing Basket Cell (PVBC) | 5530 |
| Axo-axonic Cell (AAC) | 1380 |
| Bistratified Cell (BS) | 2210 |
| Cholecystokinin-expressing Basket Cell (CCKBC) | 3600 |
| Ivy Cell (IVY) | 8810 |
| Neurogliaform Cell (NGFC) | 3580 |
| Oriens-Lacunosum Moleculare Cell (OLM) | 1640 |
| Schaffer Collateral-Associated Cell (SCA) | 400 |
| Interneuron-Specific Cell Type 1 (IS1) | 400 |
| Interneuron-Specific Cell Type 2 (IS2) | 1970 |
| Interneuron-Specific Cell Type 3 (IS3) | 1250 |
| Cornu Ammonis Region 3 Cell (CA3) | 204700 |
| Cornu Ammonis Region 2 Cell (CA2) | 40000 |
| Entorhinal Cortex Cell (EC) | 250000 |

**Figure 6.**
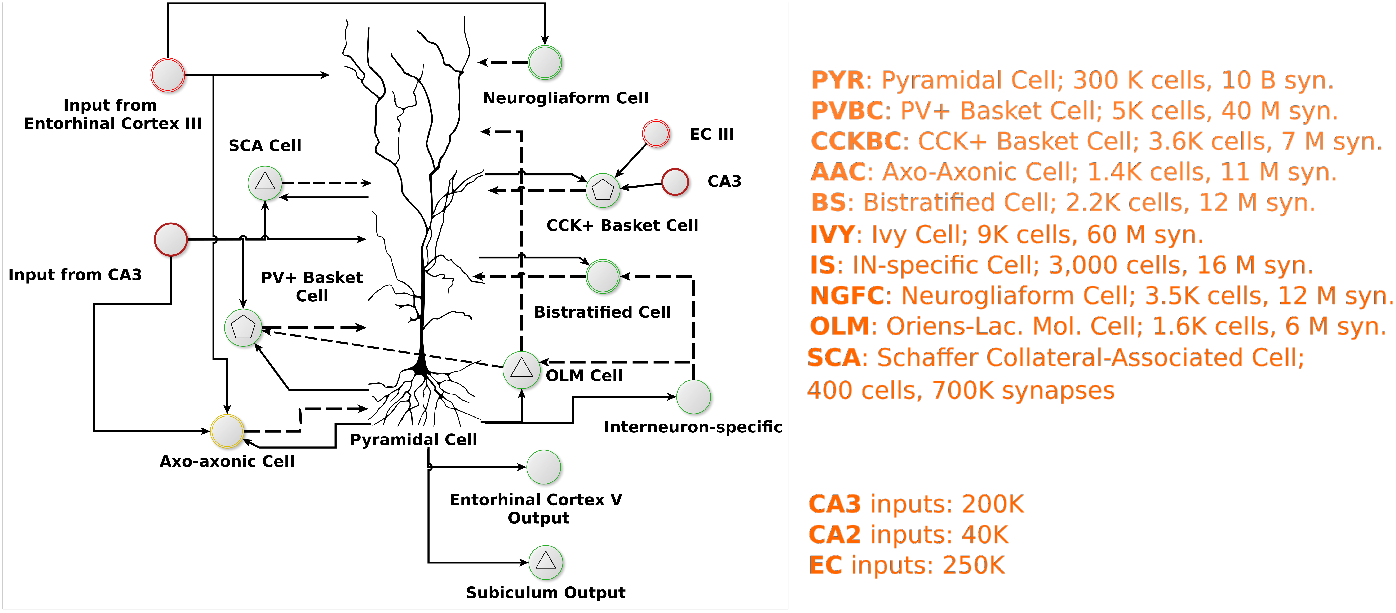
Schematic diagram of CA1 hippocampal circuit.

#### A.1.3 Synaptic noise

*livn* includes mechansims for noise generation that account for the fact that neural processes are inherently stochastic. For simpler (S)LIF models, we inject random current as an additional term. For the NEURON models, we model the stochastic nature of synaptic vesicles that are known to spontaneously release neurotransmitters even in the absence of evoked activity. Specifically, we leverage a simplified point-conductance model by Destexhe et al. (2001) that represents the currents generated by thousands of stochastically releasing synapses.

To support debugging, *livn* also includes an option for deterministic noise that is seeded to be exactly reproducible even when models are simulated on multiple workers. This simpler implementation is realized as an Ornstein-Uhlenbeck process that can be written as:

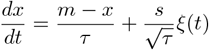

where *x* is the state variable (current amplitude), *m* is the mean value of the process (steady-state expected current), *s* is the standard deviation (square root of the variance), *τ* is the correlation time constant and *ξ*(*t*) represents white noise with zero mean and unit variance.

#### A.1.4 Electrode stimulation

To model electrode cell stimulation, we compute the induced voltage *v* for each neuron as:

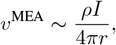

where *ρ* = 10 Ω · m is the tissue resistivity, *I* is the electrode current and *r* is the distance between the neuron and the electrode tip. For NEURON models, the resulting potential is applied using the NEURON’s built-in extracellular mechanism. For the (S)LIF and Izhikevich, the stimulus is added as additional current term, converted via Ohm’s law assuming a membrane resistance of 400*M* Ω. To avoid unrealistically high induction near *r* = 0, we limit the minimum distance between the electrode and the neuron to 5*µm*.

### A.2 Parameter tuning

All model parameters such as the ion channel conductances or synaptic projection weights are determined via constraint multiobjective optimization. In each case, we run a NSGA-II (Deb et al., 2002) black-box optimization for 10 epochs with a population size of 100 and 10 generations per epoch, drawing 100 initial parameters per search space dimension. We sort the pareto front to pick the solution with the lowest standard deviation across the objective values.

To reproduce overall plausible firing dynamics, we optimize the projection strength between each cell population (e.g. excitatory-inhibitory, excitatory-excitatory, and so forth). We define six objectives computed for *spontaneous* and/or electrode *stimulated* activity as follows:

- **Firing rate (spontaneous)** Targets a specific neuronal firing rate of 3.0 Hz as observed by (Zhang et al., 2023), minimizing the squared error between the achieved and target firing rate.
- **Active neurons ratio (spontaneous)** Aims for a 60% fraction of active neurons. The objective minimizes the squared error between the achieved and target activation ratio.
- **Inter-Spike Interval Regularity (spontaneous and stimulated)** Evaluates the regularity of firing patterns by calculating the coefficient of variation (CV) of inter-spike intervals for each neuron, then averaging across neurons. The goal is to achieve a CV close to 1.0, which represents Poisson-like firing (balanced between regular and irregular spiking).
- **Functional correlation (stimulated)** Maximizes a correlation score that measures functional connectivity between neurons, aiming for a correlation value between 0.05 and 0.4.
- **Active spiking (stimulated)** For each stimulation channel, maximizes spike counts in the early part of the trial (*t* ≤ 150ms), representing responsive activity to the stimulus.
- **Calming spiking (stimulated)** For each stimulation channel, minimizes spike counts in the later part of the trial (*t >* 150ms), ensuring temporal specificity of responses.

All the resulting optimized cell model and population projection parameters can be found in the code repository.

### A.3 Dataset generation details

The datasets for the S1-S4 system are generated by simulating a second of neural activity in response to random electrode stimuli. The input feature space consists of three parameters: the electrode channel (0-15), the time when the stimulus occurs (0 to 900 ms, ensuring a 100 ms response window before the end of the trial), and the stimulus amplitude (between 250-750 mV). Samples are drawn uniformly in each dimension, and the resulting spike trains are recorded. Each dataset is generated with and without additional synaptic noise.

For data generation for the hippocampal systems C5 and C6 rely, the input signals are converted to spike trains using a two-stage process: 1) A feature-specific encoder transforms signal samples to instantaneous firing rate maps using either:

1. Linear rate encoding: *r*(*t*) = *r*_*max*_*s*(*t*)

2. Receptive field encoding: *r*(*t*) = *r*_*max*_ exp(− ((*s*(*t*) − *c*)*/w*)^2^) (where *c* is field center, *w* is field width)

2) A Poisson spike generator converts these firing rates to spike trains with probability: *P* (*spike* | *t*) = *r*(*t*)*δt*. This approach allows for flexible modeling of diverse neural responses and can be extended to implement various tuning profiles observed in neural sensory systems.

### A.4 Dynamical response characterization

We implement a systematic framework for characterizing the computational capabilities of biophysical neural networks through mapping of their responses to diverse spatio-temporal input patterns. This Dynamical Response Characterization (DRC) system samples the high-dimensional feature space of possible neural inputs in a standardized way, enabling quantitative assessment of a network’s information processing capabilities. By systematically generating features that span several dimensions and varying temporal dynamics, frequency sensitivity, spatial selectivity, and tuning width, the system provides statistically robust coverage of theoretically relevant input patterns while minimizing redundancy in testing.

The DRC framework uses a modular architecture that separates input modalities from feature populations, allowing researchers to characterize network responses across multiple experimental paradigms while maintaining a consistent approach. Each input feature in the population can apply a specific filter function to input signals based on its position in the feature space, creating a distributed encoding system.

Using this infrastructure, *livn* allows implementation of DRC benchmarks by allowing precise control over input delivery mechanisms that mirror experimental techniques. Through a modular encoding pipeline, researchers can translate abstract feature representations into physiologically realistic spike trains for synaptic input or spatiotemporally structured light patterns for simulated optogenetic stimulation. This dual-input capability allows benchmarks to quantify computational properties under conditions that parallel both natural neural communication and experimental manipulation paradigms.

Figure 7 illustrates how temporal features are encoded and converted to spiking input. Figure 8 illustrates a multidimensional input surface of spatio-temporal input features generated with the DRC approach.

**Figure 7.**
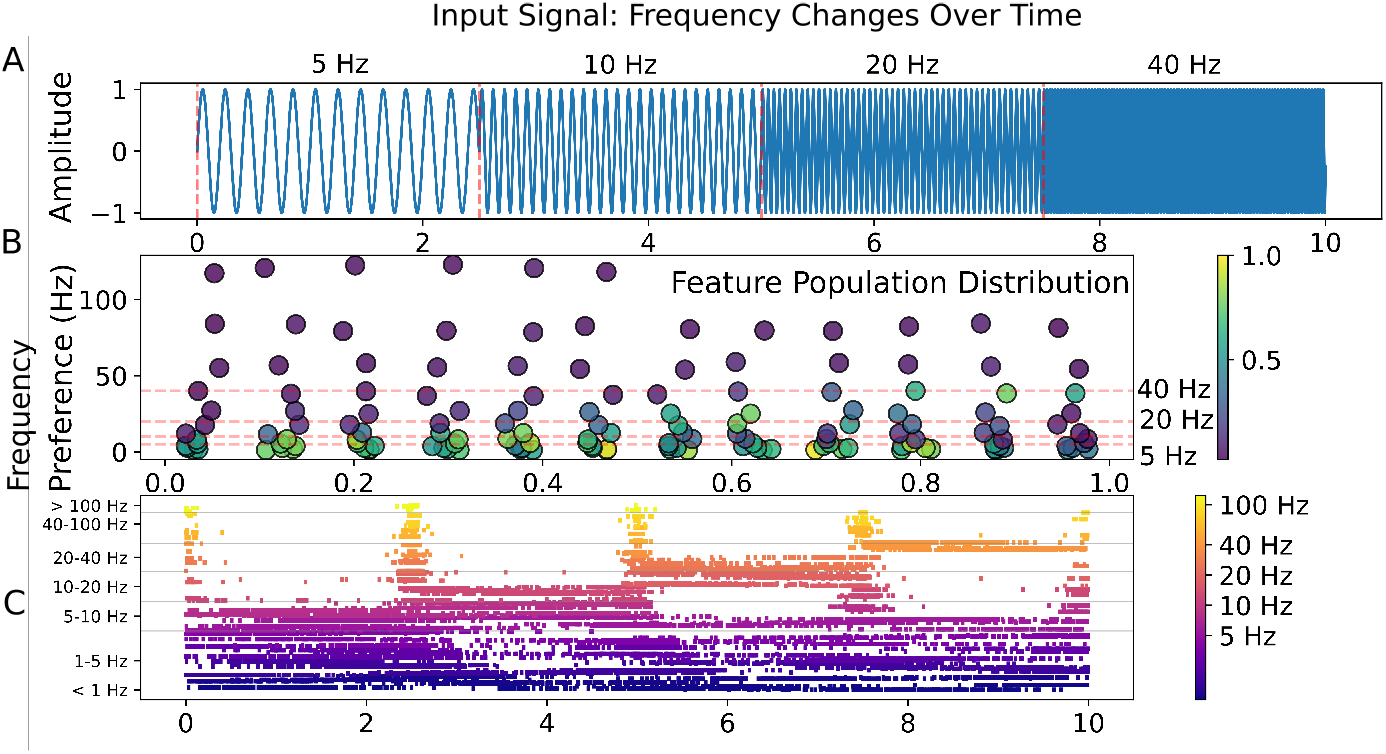
An illustration of how temporal features are encoded and converted to spiking input. A) An input signal that transitions through four distinct frequency segments (5Hz, 10Hz, 20Hz, and 40Hz), providing a systematic probe of temporal processing across the frequency spectrum. B) Distribution of feature-selective inputs in feature space, plotting their preferred temporal position against their preferred frequency. The color intensity represents the mean activation level of each feature in response to the input signal, revealing which feature-selective input units are most strongly engaged by the stimulus. C) Spike raster plot of the feature population’s response to the input signal. The distinct activation patterns across different frequency bands demonstrate the selective responses of specialized feature detectors, with each input showing highest activity when the input frequency matches its tuning preference.

**Figure 8.**
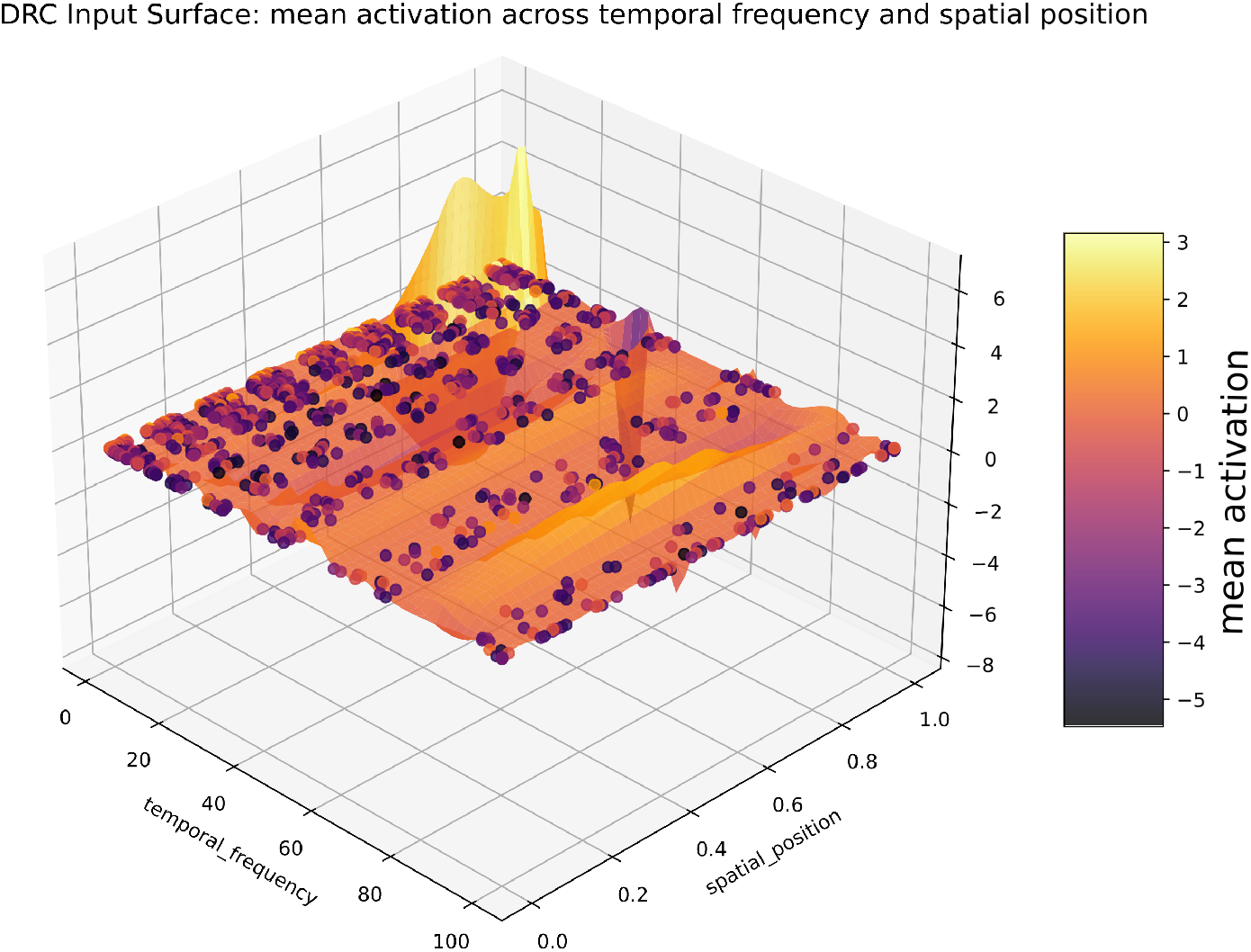
Visualization of a multidimensional input surface generated by the Dynamical Response Characterization (DRC) approach, showing mean activation levels across temporal frequency (1-100 Hz) and spatial position (0-1, normalized) dimensions. The DRC methodology generates a population of input features that act as selective filters across multiple dimensions. Each dot represents an individual feature’s position in the 2D parameter subspace (n=1250 features total), while the interpolated surface illustrates the population-level mean activation response to a test stimulus containing different frequency components (5, 10, 20, and 40 Hz) distributed across spatial and temporal regions.

## Footnotes

1 https://github.com/livn-org/livn

## Notes

### Competing Interest Statement

The authors have declared no competing interest.

